# Linear B-cell epitope prediction for *in silico* vaccine design: a performance review of methods available via command-line interface

**DOI:** 10.1101/833418

**Authors:** Kosmas A. Galanis, Katerina C. Nastou, Nikos C. Papandreou, Georgios N. Petichakis, Diomidis G. Pigis, Vassiliki A. Iconomidou

## Abstract

Linear B-cell epitope prediction research has received a steadily growing interest ever since the first method was developed in 1981. B-cell epitope identification with the help of an accurate prediction method can lead to an overall faster and cheaper vaccine design process, a crucial necessity in the covid-19 era. Consequently, several B-cell epitope prediction methods have been developed over the past few decades, but without significant success. In this study, we review the current performance and methodology of some of the most widely used linear B-cell epitope predictors which are available via a command-line interface, namely BcePred, BepiPred, ABCpred, COBEpro, SVMTriP, LBtope, and LBEEP. Additionally, we attempted to remedy performance issues of the individual methods by developing a consensus classifier, which combines the separate predictions of these methods into a single output, accelerating the epitope-based vaccine design. While the method comparison was performed with some necessary caveats and individual methods might perform much better for specialized datasets, we hope that this update in performance can aid researchers towards the choice of a predictor, for the development of biomedical applications such as designed vaccines, diagnostic kits, immunotherapeutics, immunodiagnostic tests, antibody production, and disease diagnosis and therapy.

## 1. Introduction

B-cell epitopes are regions on the surface of an antigen, where specific antibodies recognize and bind to, triggering the immune response. This interaction is at the core of the adaptive immune system, which among others is responsible for immunological memory and antigen-specific responses in vertebrates. The ability to identify these binding areas in the antigen’s sequence or structure is important for the development of synthetic vaccines [1–3], diagnostic tests [4], and immunotherapeutics [5,6], especially in the COVID-19 era. Focus on these applications through the lens of epitope discovery has gained attention over the years, especially in regard to the safety benefits of synthetic vaccine development [7].

Generally, B-cell epitopes are divided into two categories: linear (continuous) epitopes, that consist of a linear sequence of residues, and conformational (discontinuous) epitopes which consist of residues that are not contiguous in the primary protein sequence but are brought together by the folded protein structure [8]. Moreover, about 90% of B-cell epitopes have been estimated to be conformational and only about 10% to be linear [9]. Nonetheless, it has been shown that many discontinuous epitopes contain several groups of continuous residues that are also contiguous in the tertiary structure of the protein [10], making the distinction between them unclear.

All aforementioned immunological applications share the need for the discovery of all possible epitopes for any given antigen, a process called “Epitope mapping”. Although epitope mapping can be carried out using several experimental techniques [11], it is time-consuming and expensive, especially on a genomic scale. To address this problem and tap into the ever-growing data on epitopes deposited in biological databases daily, several computational methods for predicting conformational or linear B-cell epitopes have been published over the last decades [12–14] (Supplementary Table 1). Despite the relatively small percentage of linear B-cell epitopes, most methods developed over the past few years focus on their prediction. This is mainly attributed to the requirement of an antigen’s 3D structure when predicting its conformational epitopes [15]. Thus, in this review, we will discuss solely the performance of linear B-cell epitope (BCE) predictors.

Here, we review the performance of some of the most widely used linear B-cell epitope predictors currently available via a Command-Line Interface (CLI), namely BcePred [16], BepiPred [17], ABCpred [18], COBEpro [19], SVMTriP [20], LBtope [21] and LBEEP [22]. We also examine the performance of a consensus classifier combining these methods, to test whether a consensus approach can boost predictive performance [23–25]. Finally, we compare the performance of all these classifiers and the consensus method we developed against one of the most recently published BCE predictors, BepiPred-2.0 [26]. This review aims to give non-expert researchers an overview of available linear BCE predictors, as well as an update in their current performance and availability, which they can use to quickly locate and choose the appropriate tools for their research work. Moreover, we have created contemporary non-redundant datasets of linear BCEs that could aid both experimental researchers as well as bioinformaticians actively working in the field of algorithm development.

## 2. Materials and Methods

### 2.1. Selection of suitable linear B-cell epitope predictors

The first priority of this work was to gather and test as many individual predictors as possible. However, the scope of methods that were to be tested could not be limitless, and thus some criteria for their selection were applied. At first, we decided to catalogue all available B-cell epitope predictors (Supplementary Table 1). This is when we first noticed an alarming trend; where many of the online tools of the predictors that we looked up were either offline for some hours during the day or – even worse – completely unreachable. Furthermore, even when operational, most prediction servers have limitations on the amount of sequences and the workload they can process. Considering the present issues and the future problems that might arise, we decided to resort only to methods that were available as standalone software, which became our main selection criterion. The second criterion was that methods should be usable via a CLI and not only through a Graphical User Interface (GUI) and the third criterion was that each method’s way of operation should be somewhat comparable and in tune with the rest of the available predictors. Out of the many methods that have been developed through the years (Supplementary Table 1), seven were selected for testing: **BcePred**[16], **BepiPred**[17], **ABCpred**[18], **COBEpro**[19], **SVMTriP**[20], **LBtope**[21] and **LBEEP**[22]. During our study the second version of BepiPred was released, and its comparison with the rest of the methods and our decision not to utilize it in the development of the consensus method is discussed later in this article.

Once all methods were installed in a local Unix-based machine, their output was validated by comparing example sequences of the local versions of software with the corresponding online tools. Additionally, all methods used in this analysis had their threshold set on its default value except for BcePred and COBEpro (Table 1). In the case of BcePred the default threshold value of the method used, which combines the results of four different propensity scales, was decreased from 2.38 to 2. This decrease was decided after extensive testing because the default threshold value proved to be extremely high. Nevertheless, it should be noted that the new value used agreed with the default threshold currently used by both the online and the local version of the method, in contrast with the one reported in the initial publication. COBEpro on the other hand didn’t have a default threshold value, since its results are printed out in a chart where epitopic propensity is given a relative positive or negative score for each position of the query protein. The threshold value that was chosen for this method was that of four positive votes above the baseline score of zero because it yielded the best results during testing.

**Table 1.** A summary of methods, threshold values, and modifications applied to each predictor. Each predictor first had its best performing mode selected and its threshold value set to a specific value shown in the table, using the criteria described in the manuscript.

| *Predictor* | *Threshold* | *Mode* | *Threshold Type* |
| --- | --- | --- | --- |
| **BcePred** | 2 | Combined | Not Default |
| **BepiPred-1.0** | 0.35 | BepiPred | Default |
| **ABCpred** | 0.51 | 20 | Default |
| **COBEpro** | 4 | - | Not Default |
| **SVMTriP** | 0.2 | 20 | Default |
| **LBtope** | 0.6 | - | Default |
| **LBEEP** | 0.6 | Balanced | Default |

A consensus method was developed to incorporate all available methods that were selected in the first stage, and it is available upon request, due to free distribution limitations, at http://thalis.biol.uoa.gr/BCEconsensus/ as a standalone application. All sequence-based methods can be divided into two categories based on their classification approach. The first category comprises of the methods that assign an epitopic propensity score to each residue of the provided sequence. Four methods are included in it: BcePred, BepiPred, COBEpro, and LBtope. The second category comprises of the methods that classify peptides within certain length sizes as epitopes or non-epitopes, such as ABCpred, SVMTriP, and LBEEP. The two categories are summarized in Supplementary Table 2.

### 2.2. Data sets

Typically, the development of machine learning classifiers requires both a training data set and a test data set, but since all the predictors tested in this work were previously developed, only the latter was necessary. However, due to the fact that the individual training data sets for each predictor contained a significant number of overlapping sequences, gathered from a select few databases (like IEDB [27] and Bcipep [28]), their inclusion in our test data set would introduce bias in the results. So, in order to test all the different methods in an unbiased manner, the positive and negative training data sets for each method were gathered from their respective publications and webpages. As shown in Table 2, the positive training data set for the majority of predictors comprises of all available BCEs from a given database, while the negative set contains random amino acid sequences from Swiss-Prot [29]. The way the negative set of control data is constructed, changed in algorithms developed after 2012 to include only sequences from confirmed non-epitopes, as is the case for SVMTriP, LBtope, and LBEEP. This change was introduced in order to improve the ability of prediction algorithms to effectively distinguish “epitopic” from random sequences, as it had been previously proposed [30].

**Table 2.**
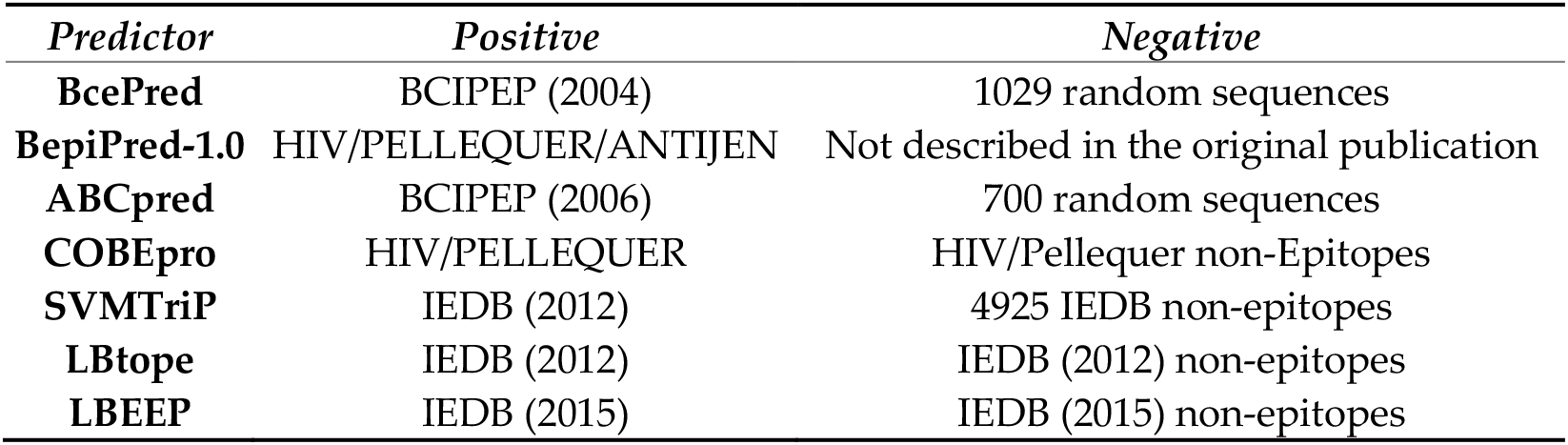
A summary of the source of positive and negative data sets for each predictor. For every predictor, a database had to be used to construct its training data sets, which comprise of a positive and a negative subset of data. In this table, we outline the database or curated data set from which each method sourced its training data set, along with the date that the data was obtained. The date could be used to determine the snapshot of the data, which could have been obtained for each predictor’s training, allowing us to determine possible overlaps of our testing data set with the relevant training data.

While developing the consensus algorithm, a new version of BepiPred was published called BepiPred-2.0 [26]. Even though the method itself wasn’t utilized in the development of the consensus method, its curated publicly available data set of linear epitopes was used as the source for this work’s data sets. This data set represents the biggest collection of linear epitope and non-epitope data used for the development of a prediction method to date, as IEDB is the largest and most frequently updated epitope database[31]. The BepiPred-2.0 data set was created by procuring from this database, all available epitopes (positive assay results) and non-epitopes (negative assay results), which were confirmed as such from two or more separate experiments. Afterwards, all peptides with a length smaller than 5 and longer than 25 residues were removed from the data set, because epitopes are rarely found outside this range [32]. Any epitopes that were found both in the positive and negative subsets were also removed. The resulting data set contains 11834 epitopes in the positive subset and 18722 non-epitopes in the negative subset. Aside from its curation, a useful feature of this data set was the mapping of all epitopes and non-epitopes on their respective parent protein sequence. This made extending each epitope to a desired length much easier.

The predictors that used IEDB as their source of epitope data are SVMTriP, LBtope, and LBEEP (Table 2). In order to produce an unbiased data set, their data sets were compared with BepiPred-2.0’s data set and all the matching peptides were removed. This resulted in our first data set, named Consensus_Redundant (Consensus_R) which comprises of 7675 epitopes and 15617 non-epitopes. Using this data set as the source, a second non-redundant data set was constructed, by clustering peptides with the online tool CD-HIT [33]. All parameters were set to default and the sequence identity cut-off was set to 0.6 or 60%, as previously done in LBEEP’s data set creation [22]. The resulting data set was named Consensus_Non_Redundant (Consensus_NR) and it includes 4286 epitopes and 5266 non-epitopes. By creating the Consensus_NR data set in this manner, we essentially made the largest non-redundant data set possible, which contained known sequences that none of the predictors had previously “seen”. Additionally, from the Consensus_NR data set a subset was extracted, containing 552 epitopes and 480 non-epitopes with a peptide length of exactly 20 amino acids, which was named Consensus_NR_exact. This subset was used to test the performance of predictors using only true epitopes and not epitope containing regions that result from the extension-truncation technique. A summary of all data sets used in this study is presented in Table 3, while the complete data sets are provided in Supplementary Table 3.

**Table 3.** A summary of test data sets utilized in this study. The counts of positive and negative subsets of data used in each of the three data sets developed for method testing is shown.

| *Data set* | *Epitopes* | *Non-Epitopes* |
| --- | --- | --- |
| **BepiPred-2.0\*** | 11814 | 18689 |
| **Consensus_R** | 7675 | 15617 |
| **Consensus_NR** | 4286 | 5266 |

\*A slightly modified version of BepiPred-2.0's data set was used, which had a few epitopes removed because their sequence of origin was shorter than 20 amino acid residues and thus the epitope couldn't be extended to the desired length.

Each data set used for testing contained peptides modified beforehand into fixed-length patterns using the technique of sequence extension and truncation, employed in previous methods [34,19,12,18]. This was done to accommodate the fixed-size input methods and thus included only their corresponding input lengths, namely 10, 12, 14, 16, 18, and 20 residues. For example, for a window size of 20, any epitopes or non-epitopes that were longer than 20 amino acids were shortened from both sides to have the desired length. Moreover, peptides with a length shorter than 20 residues were extended sideways on their parent sequence up to the desired length. The primary input size that was tested in this study was that of 20 residues for performance reasons as described in the development of the consensus algorithm. However, preliminary testing was also performed on a length of 16 residues, after analyzing the distribution of epitope lengths in the BepiPred-2.0 data set (Supplementary Figure 1). The mean peptide length of the data set was about 14 and the median value 15, which coincides with previous research on the characteristics of epitopes [32].

The workflow used to create the non-redundant data sets is shown in Supplementary File 1 (Figure 3) and all data sets referenced in this section can be downloaded from this web page http://thalis.biol.uoa.gr/BCEconsensus/.

### 2.3. Performance measures

To evaluate a classifier’s performance both threshold dependent and independent metrics are used. The main threshold independent metric used in such cases is the AUC of the ROC curve. This metric was suggested as the preferred metric for benchmarking epitope prediction performance at a workshop by Greenbaum *et al.* [30] and thus, it grew to become a standard in the BCE prediction field. However, because all the predictors that we examined were already fully developed and their optimal thresholds set, it didn’t make sense to use such a metric in our testing, since no model training was performed. For that reason, only threshold dependent metrics were employed, namely Sensitivity (SN), Specificity (SP), Accuracy (ACC), and Matthew’s Correlation Coefficient (MCC). Out of these metrics, significant attention was given to MCC, since it is generally regarded as the best performance metric for binary classifiers [35,36]. The coefficient’s value can range from −1 to +1, where the maximum value represents a perfect prediction and the minimum a total disagreement between predictions and observations. When the coefficient’s value is zero it indicates a prediction that is no better than random. Aside from the known value in accessing performance utilizing the MCC and accuracy metrics, regarding the other metrics, more importance was attached to sensitivity rather than specificity. Sensitivity indicates how effectively a predictive method manages to successfully locate areas that are actual epitopes, in contrast to specificity, which measures how effectively a predictive method manages to locate the sites that are not epitopes. In this study, the correctly predicted epitopes or “epitopic” residues were considered True Positive (TP), whereas the correctly predicted non-epitopes or “non-epitopic” residues were characterized as True Negative (TN). Conversely, the respective false predictions were defined as False Positive (FP) and False Negative (FN), respectively.

## 3. Results and Discussion

As mentioned in the *Methods* section, two approaches are followed to evaluate all predictions made by the consensus algorithm. In the first approach results from all methods are incorporated in the consensus method — both those predicting in a “per residue” and in a “per peptide” manner — while in the second approach the consensus prediction only utilizes the “per residue” methods. Two different versions of the consensus algorithm were created in the “per peptide” mode, as seen in Table 4; one which includes all predictors and one which utilizes all of them except LBEEP. This was done after noticing that LBEEP performs much worse, compared to the rest of the predictors. This performance issue can be mainly attributed to the fact that the optimal prediction window of 5-15 residues for LBEEP is different than the 20-residue length that was used for our testing purposes (Supplementary Table 2).

**Table 4.** Performance of all predictors in “per peptide” mode. The methods are tested against the Consensus_NR data set.

| *Predictor* | *SN%* | *SP%* | *ACC%* | *MCC* |
| --- | --- | --- | --- | --- |
| **Consensus_noLBEEP** | 48.39 | 58.81 | 54.14 | **0.0721** |
| **Consensus_ALL** | 27.15 | 78.73 | 55.59 | 0.0687 |
| **BcePred** | 22.21 | 79.85 | 53.99 | 0.0251 |
| **ABCpred** | 66.44 | 36.9 | 50.16 | 0.0348 |
| **LBtope** | 45.91 | 58.94 | 53.1 | 0.0488 |
| **BepiPred-1.0** | 49.95 | 57.84 | 54.3 | **0.0778** |
| **COBEpro** | 58.63 | 45.67 | 51.49 | 0.0431 |
| **SVMTriP** | 16.21 | 85.87 | 54.62 | 0.0290 |
| **LBEEP** | 19.06 | 80.12 | 52.72 | -0.0103 |

The evaluation of the predictors’ performance was done primarily by measuring their MCC values, while secondary importance was assigned to achieving higher accuracy, and sensitivity. Sensitivity was considered more important than specificity for this particular application since a BCE predictor’s primary goal is to find possible BCEs in unknown sequences. Naturally, sensitivity and specificity are not mutually exclusive entities, yet in this study optimal sensitivity is preferred to optimal specificity. For further testing results please refer to Supplementary File 2.

### 3.1. Performance of all predictors on Consensus_NR

The results regarding the “per peptide” approach (Table 4) show that the highest MCC value was achieved by the BepiPred method with 0.0778, followed by our Consensus_NoLBEEP algorithm — the one without LBEEP — that achieved an MCC of 0.0721. Moreover, LBEEP had the lowest MCC (−0.0103), while BcePred and SVMTriP also scored low (0.0251 and 0.0290, respectively). The highest accuracy was achieved by our Consensus_ALL method with 55.59%, which was marginally better than those of SVMTriP and BcePred. SVMTriP had the best specificity out of all the methods (85.87%), followed by LBEEP and BcePred. Additionally, the ABCpred method achieved the greatest sensitivity with 66.44%, and COBEpro achieved the second highest with 58.63%. The Consensus_NoLBEEP algorithm achieved values close to the best for both MCC and accuracy, and also had a relatively improved MCC and a significantly increased sensitivity compared to its first version.

In the case of the “per residue” approach (Table 5), the consensus method (Consensus_RES) achieved the best MCC with 0.489, while BepiPred scored marginally worse with 0.0488. The same pattern was also observed for accuracy, where the Consensus_RES method scored 53.04% and BepiPred 52.88%. The greatest sensitivity was achieved by COBEpro with 49.27%, while BepiPred was again second best with 48.12%. The worst performance regarding MCC was attained by BcePred and COBEpro with scores of 0.0154 and 0.0175 respectively. Overall, despite the slight improvement in MCC and accuracy, the performance of the consensus algorithm was not significantly better in any of the statistical measures examined in the second part of the results.

**Table 5.** Performance of “per residue” predictors. The methods are tested against the Consensus_NR data set.

| *Predictor* | *SN%* | *SP%* | *ACC%* | *MCC* |
| --- | --- | --- | --- | --- |
| **Consensus_RES** | 46.64 | 58.24 | 53.04 | 0.0489 |
| **BcePred** | 29.18 | 72.21 | 52.9 | 0.0154 |
| **LBtope** | 45.56 | 57.47 | 52.13 | 0.0304 |
| **BepiPred-1.0** | 48.12 | 56.76 | 52.88 | 0.0488 |
| **COBEpro** | 49.27 | 52.49 | 51.05 | 0.0175 |

When comparing the results of the two approaches only minor differences in performance are observed between the two modes of prediction for the four “per residue” methods. Generally, we notice a slight decrease in MCC from a maximum of 0.0778 in the first approach to a maximum of 0.0489 in the second, while accuracy is comparatively worse on average. Out of the “per residue” methods, BepiPred comes on top in both approaches in MCC and accuracy. The Bcepred method appears to perform relatively worse than the rest in both groups with the lowest MCC in both cases, whereas the COBEpro method performs relatively better in its “per peptide” iteration, with an average MCC score in the first part but a poor score in the second segment of the results. Moreover, in both approaches, our consensus algorithm doesn’t significantly outperform the rest of the predictors and only achieves a performance that is quite similar to that of BepiPred.

In summary, we observe that in all cases: MCC values are less than 0.1, accuracy is ranging from 50% to 55%, there are relatively high specificity values in certain cases such as SVMTriP and BcePred, and sensitivity values are low. Aside from our consensus methods, the best performers were LBtope and BepiPred and the worst ABCpred and LBEEP, which also displayed the lowest MCC scores.

Using the Consensus_NR data set we implemented many iterations of the consensus method utilizing many different method combinations, in order to find the optimum. As expected, LBEEP’s presence undermined the consensus predictor’s performance and it was therefore omitted from the final version (Consensus_NoLBEEP) and any further testing in the 20-residue window size. It was also observed that ABCpred overestimated the presence of epitopes in their respective peptides, which led to reduced accuracy and increased sensitivity. Nevertheless, it remained part of the final consensus algorithm to improve its overall sensitivity.

At this point, it should be noted that LBEEP was also tested on a peptide length of 14-residues since the method was reported to perform optimally when a window size between 5-15 residues is used for prediction. Results showed that the method indeed performs better at this window size, but it is still marginally better than a random prediction according to its MCC (Supplementary Table 4-A). Even though, the results were better for LBEEP the rest of the methods either cannot be used at that window size or perform way worse than what we had already seen and so we opted to not use the 14-residue window any further.

### 3.2. Overall method performance and comparison with BepiPred-2.0

The performance of the linear B-cell epitope predictors examined was found to be poor in the data sets and window sizes used during testing (Figure 1).

**Figure 1.**
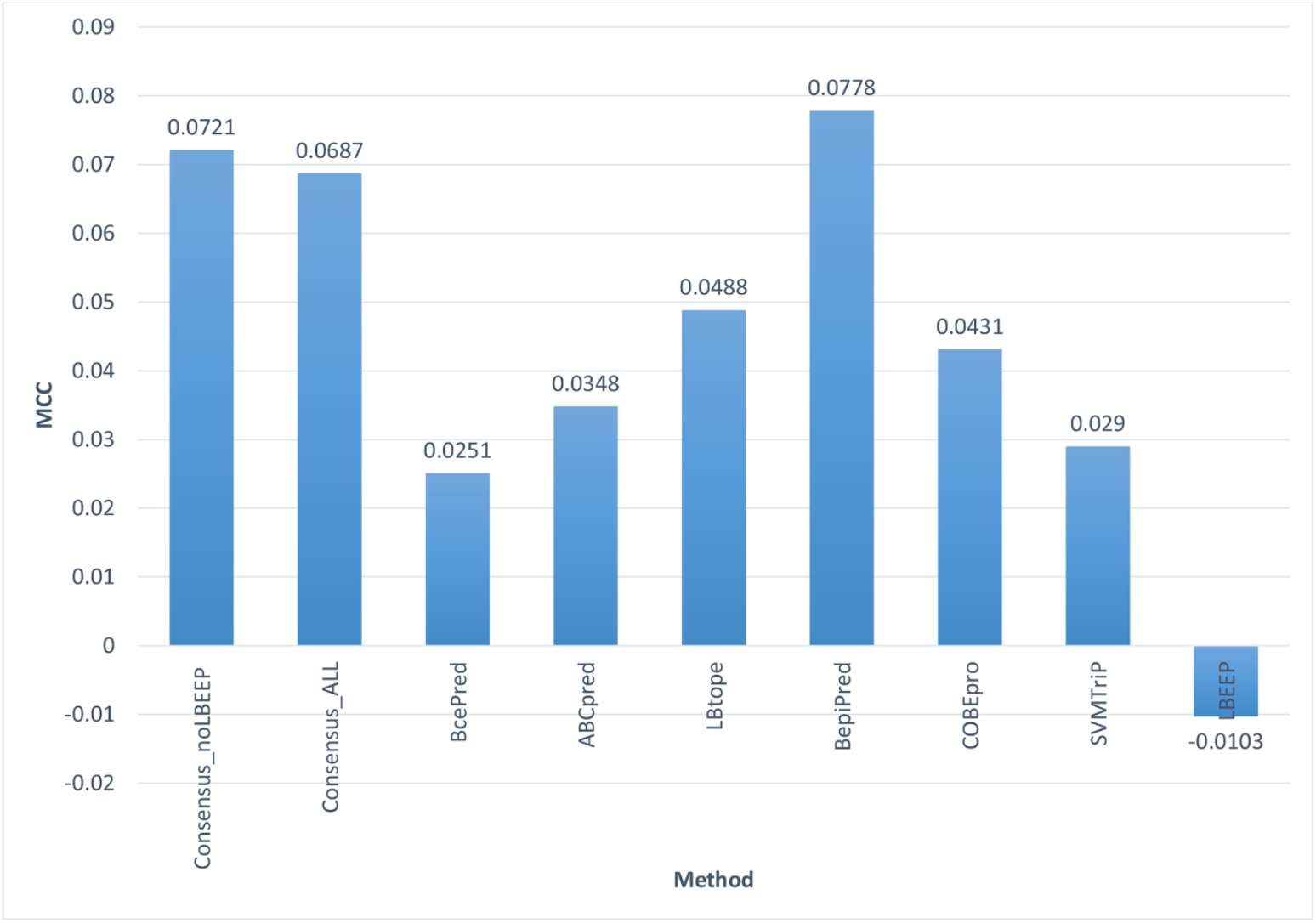
MCC values achieved by all methods tested on the Consensus_NR data set at 20 amino acid residues in “per peptide” mode. The vertical axis represents the MCC value for all the methods and the horizontal axis the names of these methods. The best MCC is achieved by the BepiPred method, followed closely by our Consensus methods, while the worst performers are the LBEEP, SVMTriP, and BcePred methods.

Additionally, despite our optimization, our consensus method performed only marginally better than the rest of the methods, thus nullifying its usefulness. We believe that the problems which may explain these results can be divided into two categories; those concerning the individual methods and those of the consensus approach.

The first problem regarding the prediction methods is that the epitope data used to train and test them, and as a result, the methods themselves are outdated. This probably is what caused their significantly reduced performance in our contemporary and considerably larger set of data. Furthermore, the general difficulty of creating a relatively reliable sequence-based predictor is well known, in contrast with those available for example in the prognosis of T-cell epitopes [37]. This is mainly due to the 3D nature of all B-cell epitopes, which consist of seemingly unrelated residue patterns of the antigen. Their emergence is also subject to multiple factors, such as antigen concentration and the type of chemical test [30].

In our attempt to create a consensus predictor, the first problem we encountered was the different modes of operation of the individual prediction methods, namely their distinction into “per peptide” and “per residue” predictors. To effectively compare the two modes, “per residue” predictor outputs were converted to “per peptide”, by using a percentage cut-off to classify peptides as epitopes and non-epitopes. This, however, is not their intended operation mode, which certainly influences the performance of these methods and thus the performance of the consensus method.

Another obstacle in this effort was time and complexity. The prediction and evaluation process for all possible windows (10, 12, 14, 16, 18 and 20) is very time-consuming. This also had to be performed for as many predictors as possible to make the consensus classifier more effective, leading to a significant increase in software development complexity as the number of incorporated predictors grew. In addition, accurate assessment of the viability of such an effort is very difficult, due to the inability to accurately compare them beforehand using the results presented in the corresponding publications, as there is no single set of evaluation data or metrics [12]. Finally, there was a lack of variety in the methods utilized in our selected predictors, where most of them were based on SVM models, which may have negatively affected the performance of our consensus predictor [38].

When comparing all of the methods we tested, with some of the newer methods such as BepiPred-2.0 and iBCE-EL, which were tested on large non-redundant data sets much like the ones we used, their reported superiority is apparent. Out of the two, BepiPred-2.0 was released during the initial part of testing in our research, and as such, it was a likely candidate for our consensus method. However, after observing the poor performance of all the different methods tested against its data set, we decided to not include it in our consensus approach, but simply to use it as a reference for what a modern predictor can achieve versus the older ones. Unlike its predecessor, BepiPred-1.0, and most other sequence-based predictors, BepiPred-2.0 is trained exclusively on epitope data derived from antigen-antibody crystal structure complexes obtained from the Protein Data Bank [39]. This was done in order to combat the generally poor performance of predictors, which can be partly attributed to poorly annotated and noisy training data, in comparison with data derived from crystal structures which is presumed to be of higher quality and indeed resulted in a significantly improved predictive power [26]. From these complexes, all antigenic residues close enough to their respective antibody were gathered. These residues became the positive subset of the training data set, while the negative subset was constructed from randomly selected non-epitope residues.

While, BepiPred-2.0 was trained using epitope data derived only from 3D structures, its performance on linear BCEs was also benchmarked on one such data set. We compared the performance of BepiPred-2.0 against our Consensus_noLBEEP predictor using the Consensus_NR dataset at a window size of 20 amino acid residues. When compared to our consensus method, BepiPred-2.0 has a similar performance in accuracy and MCC, but exhibits higher sensitivity and lower specificity, as shown in the comparison performed in Table 6. However, the results for both methods are far from optimal, and a lot of work still remains to be done in order to create a predictor that will perform optimally during linear BCE detection.

**Table 6.**
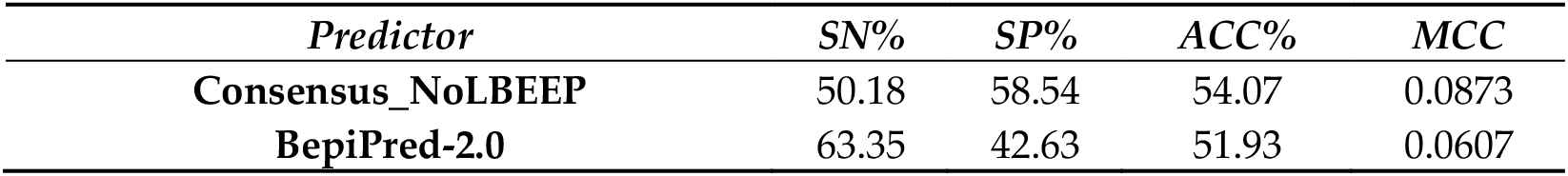
Comparison of the performance of our consensus predictor and BepiPred-2.0 against the Consensus_NR data set.

| *Predictor* | *SN%* | *SP%* | *ACC%* | *MCC* |
| --- | --- | --- | --- | --- |
| **Consensus_NoLBEEP** | 50.18 | 58.54 | 54.07 | 0.0873 |
| **BepiPred-2.0** | 63.35 | 42.63 | 51.93 | 0.0607 |

## 4. Conclusions

In summary, in this paper, we independently evaluated the performance of seven of the most popular linear B-cell epitope predictors on the largest unbiased data set possible. In the process, we also presented the course of design, development, and evaluation of a consensus prediction algorithm for linear B-cell epitopes. The performance of all predictors, except for LBEEP on whom testing was exploratory, was found marginally better than random classification. Additionally, our Consensus classifier failed to significantly outperform its constituent methods. While the method comparison was performed with some necessary compromises, we believe that this update in performance can help to better inform researchers that wish to consult some of these tools to facilitate and direct their research. Instead, we should also like to suggest that researchers opt for some of the newer predictors referenced in this work, like BepiPred-2.0. Also, due to the apparent difficulty of constructing an accurate general-purpose linear BCE predictor, we believe that software development should instead be focused on the creation of more specialized predictors for specific antigenic systems, such as known viruses or viral families of high interest, like the SARS-CoV-2 virus or other coronaviruses. This could lead to optimization in the feature selection process during classifier training and better predictive performance within that limited scope.

## Supplementary Materials

Supplementary Table 1: General information on Linear B-cell epitope predictors and supplementary introductory. Supplementary Table 2: Supplementary information on the materials and methods used and a table of input window sizes along with the prediction approach of each method. Supplementary Table 3: Evaluation Datasets. Supplementary Table 4: A) LBEEP’s performance on peptides of 14-amino acid residues. B) LBEEP’s performance using 3 different models for the 20 amino acid residue peptides of the Consensus_NR data set. Supplementary File 1: Supplementary Figure 2 showing the workflow of the consensus method and Supplementary Figure 3 showing the workflow of the creation of the non-redundant data sets. Supplementary File 2: Supplementary results and discussion. Supplementary Figure 1: Distribution of epitope lengths in the Bepipred 2.0 data set.

## Acknowledgments

We thank the National and Kapodistrian University of Athens for use of premises and equipment.

## Author Contributions

Study design: K.C.N., N.C.P., D.G.P., V.A.I.; Conceptualization: K.C.N., N.C.P., V.A.I.; Retrieval of standalones and installation: K.A.G., GNP, K.C.N.; Development of consensus method: K.A.G.; Data set acquisition and curation: K.A.G., K.C.N.; Writing - original draft: K.A.G.; Writing - review and editing: K.C.N., K.A.G., N.C.P., GNP, D.G.P., V.A.I.; Supervision: V.A.I. The manuscript was written through contributions of all authors. All authors have given approval to the final version of the manuscript.

## Abbreviations

BCE: Linear B-cell epitope
CLI: Command Line Interface
GUI: Graphical User Interface
AUC: Area Under Curve
ROC: Receiver Operating Characteristic
MCC: Matthews Correlation Coefficient
IEDB: Immune Epitope Data Base
Consensus_R: Consensus_Redundant
Consensus_NR: Consensus_Non_Redundant
SN: Sensitivity
SP: Specificity
ACC: Accuracy
TP: True Positive
TN: True Negative
FP: False Positive
FN: False Negative

## References

1. Hughes, E.E.; Gilleland, H.E.Jr. Ability of synthetic peptides representing epitopes of outer membrane protein F of Pseudomonas aeruginosa to afford protection against P. aeruginosa infection in a murine acute pneumonia model. Vaccine 1995, 13, 1750–1753.

2. Tam, J.P.; Lu Y.A. Vaccine engineering: enhancement of immunogenicity of synthetic peptide vaccines related to hepatitis in chemically defined models consisting of T- and B-cell epitopes. Proc Natl Acad Sci U S A 1989, 86, 9084–9088.

3. Russi, R.C.; Bourdin, E.; Garcia, M.I.; Veaute, C.M.I. In silico prediction of T- and B-cell epitopes in PmpD: First step towards to the design of a Chlamydia trachomatis vaccine. Biomed J 2018, 41, 109–117.

4. Schellekens, G.A.; Visser, H.; de Jong, B.A.; van den Hoogen, F.H.; Hazes, J.M.; Breedveld, F.C.; van Venrooij, W.J. The diagnostic properties of rheumatoid arthritis antibodies recognizing a cyclic citrullinated peptide. Arthritis Rheum 2000, 43, 155–163.

5. Chirino, A.J.; Ary, M.L.; Marshall, S.A. Minimizing the immunogenicity of protein therapeutics. Drug Discov Today 2004, 9, 82–90.

6. Shirai, H., Prades, C., Vita, R., Marcatili, P., Popovic, B., Xu, J., Overington, J.P., Hirayama, K., Soga, S., Tsunoyama, K., Clark, D., Lefranc, M.P., Ikeda, K. Antibody informatics for drug discovery. Biochim Biophys Acta 2014, 1844, 2002–2015.

7. Nardin, E.H., Calvo-Calle, J.M., Oliveira, G.A., Nussenzweig, R.S., Schneider, M., Tiercy, J.M., Loutan, L., Hochstrasser, D., Rose, K. A totally synthetic polyoxime malaria vaccine containing Plasmodium falciparum B cell and universal T cell epitopes elicits immune responses in volunteers of diverse HLA types. J Immunol 2001, 166, 481–489.

8. Barlow, D.J., Edwards, M.S., Thornton, J.M. Continuous and discontinuous protein antigenic determinants. Nature 1986, 322, 747–748.

9. Pellequer, J.L., Westhof, E., Van Regenmortel, M.H. Predicting location of continuous epitopes in proteins from their primary structures. Methods Enzymol 1991, 203, 176–201.

10. Van Regenmortel, M.H. Immunoinformatics may lead to a reappraisal of the nature of B cell epitopes and of the feasibility of synthetic peptide vaccines. J Mol Recognit 2006, 19, 183–187.

11. Reineke, U., Schutkowski, M. Epitope mapping protocols. Preface. Methods Mol Biol 2009, 524, v–vi.

12. El-Manzalawy, Y., Honavar, V. Recent advances in B-cell epitope prediction methods. Immunome Res 2010, 6 Suppl 2, S2.

13. Sanchez-Trincado, J.L., Gomez-Perosanz, M., Reche, P.A. Fundamentals and Methods for T- and B-Cell Epitope Prediction. J Immunol Res 2017, 2017, 2680160.

14. Tomar, N., De, R.K. Immunoinformatics: a brief review. Methods Mol Biol 2014, 1184, 23–55.

15. Flower, D.R. Immunoinformatics. Predicting immunogenicity in silico. Preface. Methods Mol Biol 2007, 409, v–vi

16. Saha, S., Raghava, G.P.S. BcePred: Prediction of Continuous B-Cell Epitopes in Antigenic Sequences Using Physico-chemical Properties. In, Berlin, Heidelberg, Artificial Immune Systems 2004, Springer Berlin Heidelberg, pp 197–204

17. Larsen, J.E., Lund, O., Nielsen, M. Improved method for predicting linear B-cell epitopes. Immunome Res2006, 2, 2.

18. Saha, S., Raghava, G.P. Prediction of continuous B-cell epitopes in an antigen using recurrent neural network. Proteins 2006, 65, 40–48.

19. Sweredoski, M.J., Baldi, P. COBEpro: a novel system for predicting continuous B-cell epitopes. Protein Eng Des Sel 2009, 22, 113–120.

20. Yao, B., Zhang, L., Liang, S., Zhang, C. SVMTriP: A Method to Predict Antigenic Epitopes Using Support Vector Machine to Integrate Tri-Peptide Similarity and Propensity. PLOS ONE 2012, 7, e45152.

21. Singh, H., Ansari, H.R., Raghava, G.P. Improved method for linear B-cell epitope prediction using antigen’s primary sequence. PloS one 2013, 8, e62216.

22. Saravanan, V., Gautham, N. Harnessing Computational Biology for Exact Linear B-Cell Epitope Prediction: A Novel Amino Acid Composition-Based Feature Descriptor. OMICS 2015, 19, 648–658.

23. Tsolis, A.C., Papandreou, N.C., Iconomidou, V.A., Hamodrakas, S.J. A consensus method for the prediction of ’aggregation-prone’ peptides in globular proteins. PloS one 2013, 8, e54175.

24. Ji, C., Ma, S. Combinations of weak classifiers. IEEE Trans Neural Netw 1997, 8, 32–42.

25. Moutaftsi, M., Peters, B., Pasquetto, V., Tscharke, D.C., Sidney, J., Bui, H.H., Grey, H., Sette, A. A consensus epitope prediction approach identifies the breadth of murine T(CD8+)-cell responses to vaccinia virus. Nat Biotechnol 2006, 24, 817–819.

26. Jespersen, M.C., Peters, B., Nielsen, M., Marcatili, P. BepiPred-2.0: improving sequence-based B-cell epitope prediction using conformational epitopes. Nucleic Acids Res 2017, 45, W24–W29.

27. Vita, R., Zarebski, L., Greenbaum, J.A., Emami, H., Hoof, I., Salimi, N., Damle, R., Sette, A., Peters, B. The immune epitope database 2.0. Nucleic Acids Res 2010, 38, D854–862.

28. Saha, S., Bhasin, M., Raghava, G.P. Bcipep: a database of B-cell epitopes. BMC Genomics 2005, 6, 79.

29. The UniProt C. UniProt: the universal protein knowledgebase. Nucleic Acids Res 2017, 45, D158–D169.

30. Greenbaum, J.A., Andersen, P.H., Blythe, M., Bui, H.H., Cachau, R.E., Crowe, J., Davies, M., Kolaskar, A.S., Lund, O., Morrison, S., Mumey, B., Ofran, Y., Pellequer, J.L., Pinilla, C., Ponomarenko, J.V., Raghava, G.P., van Regenmortel, M.H., Roggen, E.L., Sette, A., Schlessinger, A., Sollner, J., Zand, M., Peters, B. Towards a consensus on datasets and evaluation metrics for developing B-cell epitope prediction tools. J Mol Recognit 2007, 20, 75–82.

31. Vita, R., Overton, J.A., Greenbaum, J.A., Ponomarenko, J., Clark, J.D., Cantrell, J.R., Wheeler, D.K., Gabbard, J.L., Hix, D., Sette, A., Peters, B. The immune epitope database (IEDB) 3.0. Nucleic Acids Res 2015, 43, D405–412.

32. Kringelum JV, Nielsen M, Padkjaer SB, Lund, O. Structural analysis of B-cell epitopes in antibody:protein complexes. Mol Immunol 2013, 53, 24–34.

33. Huang, Y., Niu, B., Gao, Y., Fu, L., Li, W. CD-HIT Suite: a web server for clustering and comparing biological sequences. Bioinformatics 2010, 26, 680–682.

34. Lian, Y., Huang, Z.C., Ge, M., Pan, X.M. An Improved Method for Predicting Linear B-cell Epitope Using Deep Maxout Networks. Biomed Environ Sci 2015, 28, 460–463.

35. Chicco, D. Ten quick tips for machine learning in computational biology. BioData Min 2017, 10, 35.

36. Boughorbel, S., Jarray, F., El-Anbari, M. Optimal classifier for imbalanced data using Matthews Correlation Coefficient metric. PloS one 2017, 12, e0177678.

37. Konstantinou, G.N. T-Cell Epitope Prediction. Methods Mol Biol. 2017, 1592, 211–222.

38. Flower, D.R. Immunoinformatics and the in silico prediction of immunogenicity. An introduction. Methods Mol Biol 2007, 409, 1–15.

39. Berman, H.M., Westbrook, J., Feng, Z., Gilliland, G., Bhat, T.N., Weissig, H., Shindyalov, I.N., Bourne, P.E. The Protein Data Bank. Nucleic Acids Res. 2000, 28, 235–242.

