## Supplementary figures and images for "Linear B-cell epitope prediction for *in silico* vaccine design: a performance review of methods available via command-line interface"

### Supplementary Figure 1

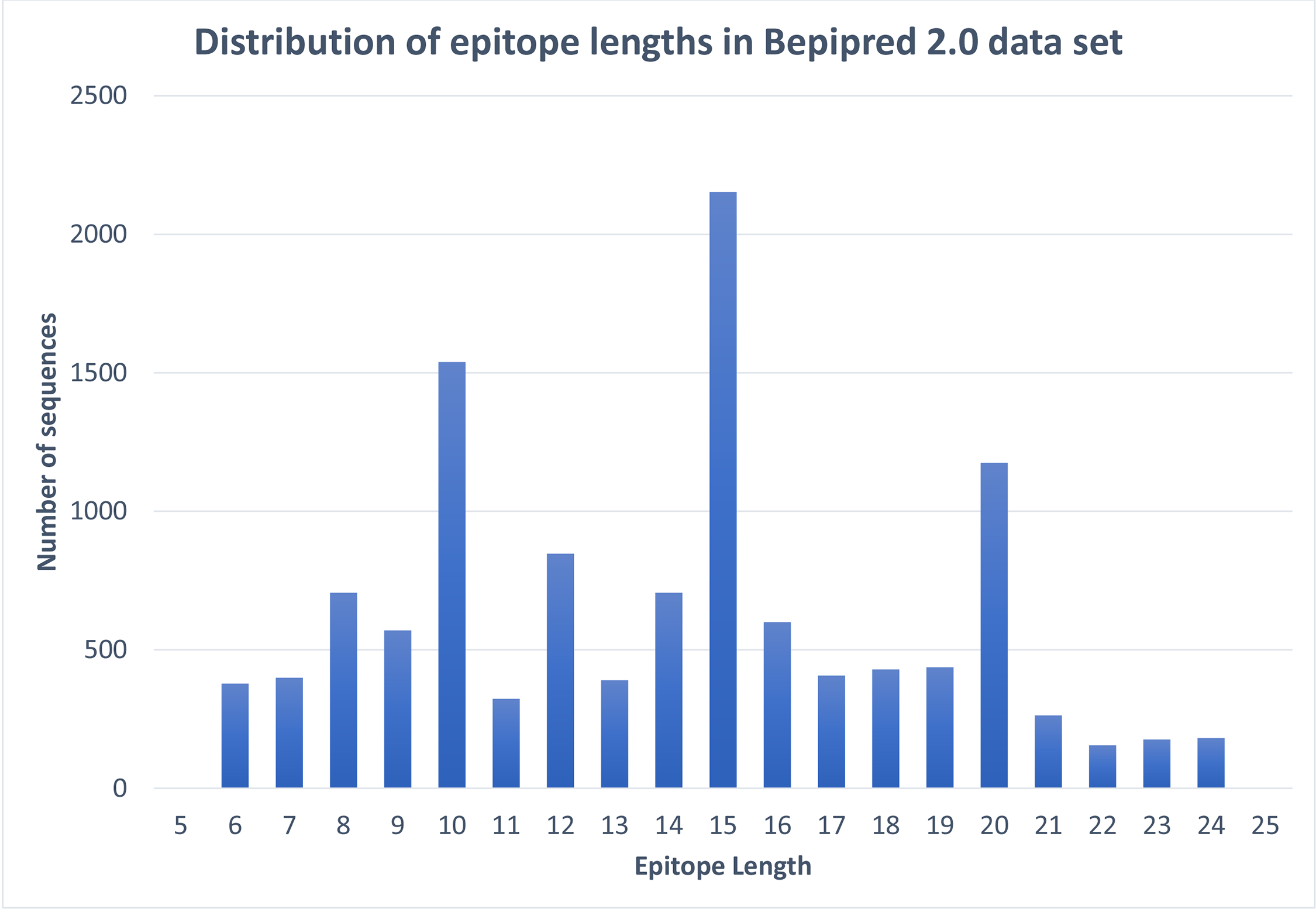
