## Supplementary File 1 for "Linear B-cell epitope prediction for *in silico* vaccine design: a performance review of methods available via command-line interface"

*Supplementary Figures*

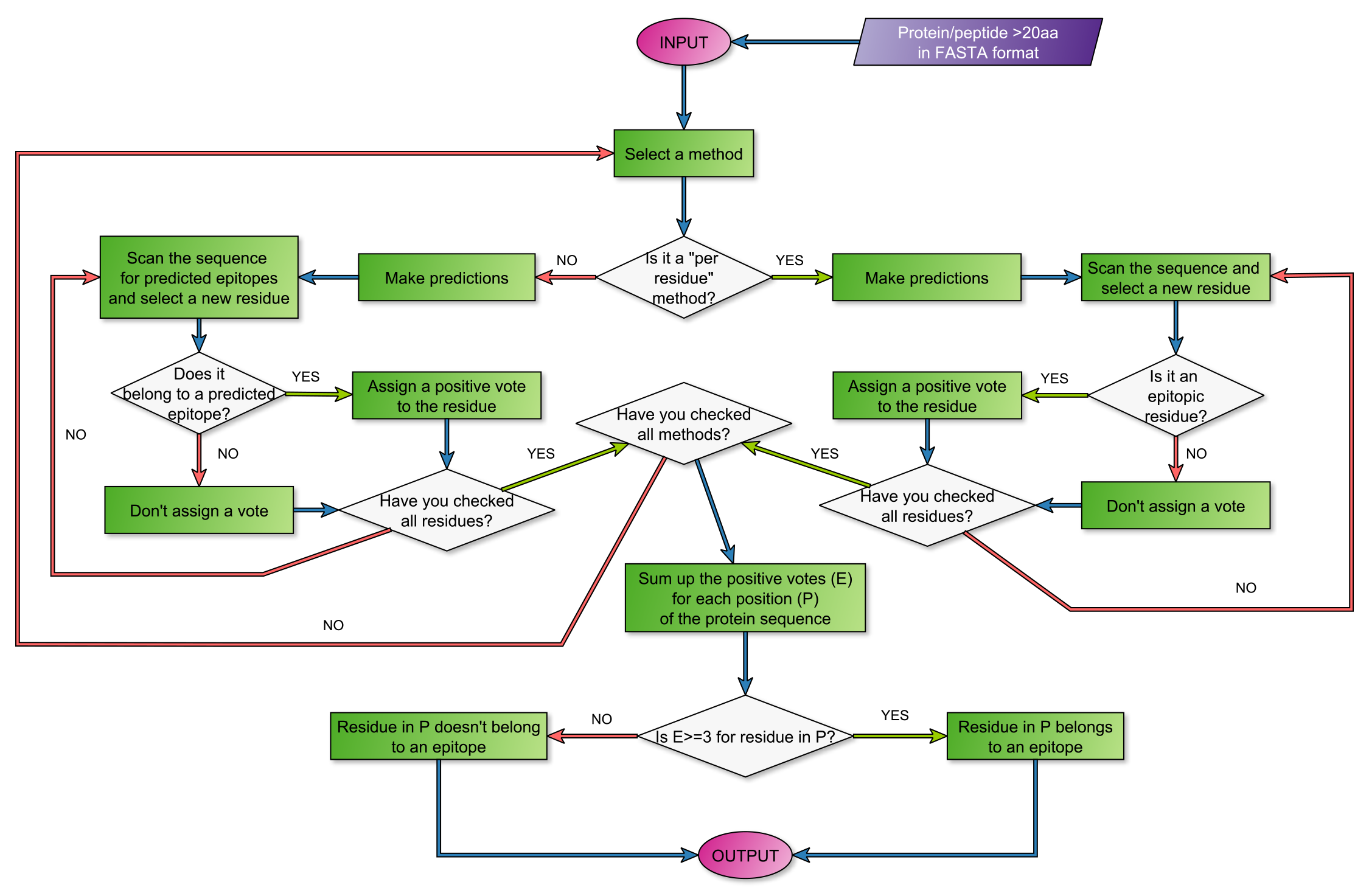

**Scheme S2.** Workflow of the consensus method. In this flowchart we present an overview of the workflow of our consensus method. The method accepts a single protein sequence in FASTA format, that is longer than 20 amino acid residues, as input. The given sequence is then passed on to every method that is part of the consensus method for prediction, namely ABCpred, BcePred, BepiPred-1.0, SVMTriP, LBtope and COBEpro. Afterwards, the results of those predictions are processed differently, depending on whether the predictor that made the prediction is a “per residue” or a “per peptide” method. The “per residue” methods, as their name implies, classify the residues of any given protein sequence as belonging to an epitope or not. Thus, for these methods our consensus method simply scans the input sequence along with its predictions and assigns a positive vote to each position of the input sequence where a residue has been classified as epitopic. On the other hand, the “per peptide” methods classify stretches of the given protein as epitopes or non-epitopes. Our consensus method, in a manner similar to that of the “per peptide” methods, checks the positions of all the residues that have been predicted as parts of an epitope and assigns a positive vote to each one of them. Finally, the positive votes for each position are summed and if the corresponding residues have a score equal or greater than 3, then they are classified as belonging to an epitope with the letter “E”, otherwise they remain unmarked. .

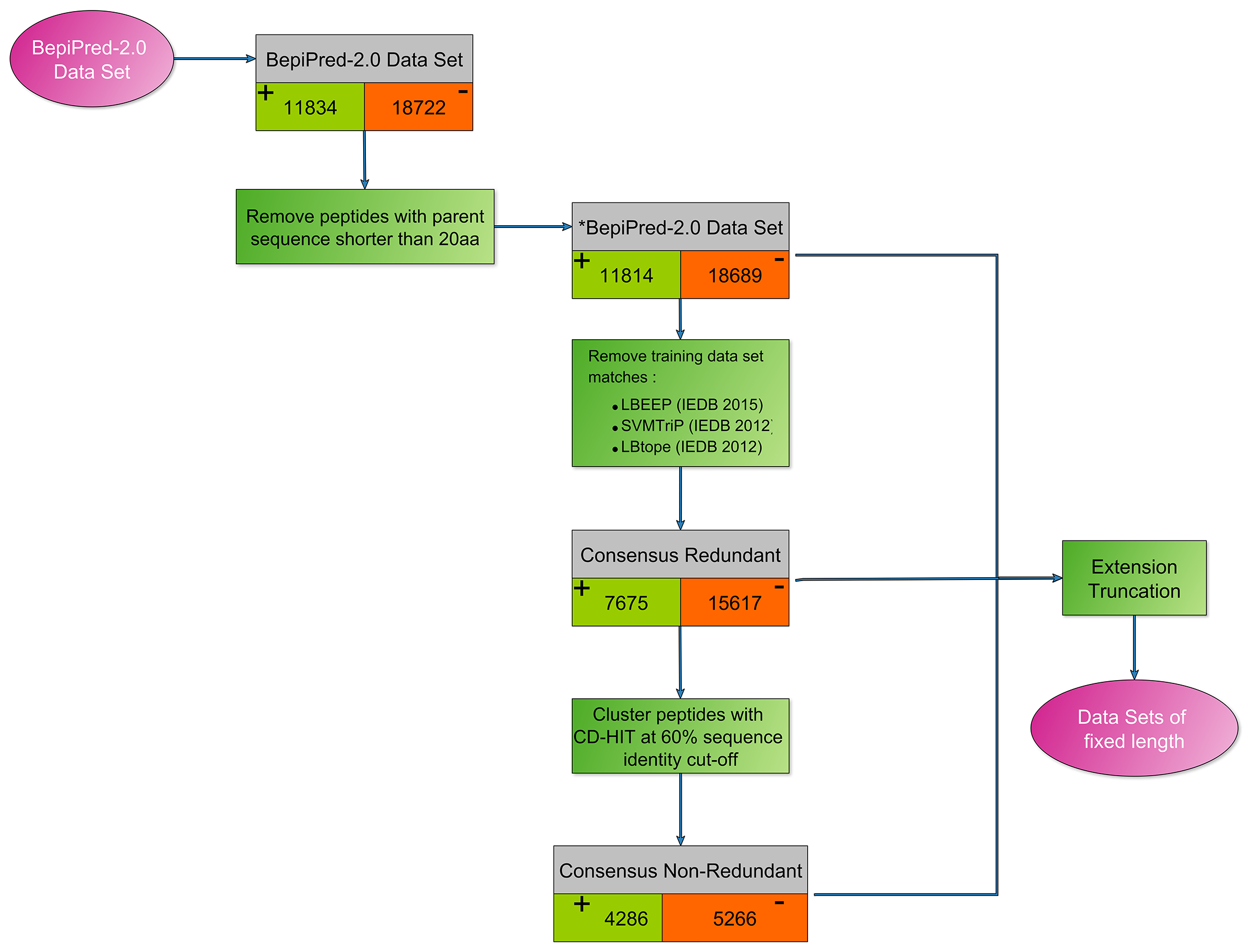

**Figure S3.** Workflow depicting the creation of the non-redundant data sets. All data sets used in the testing of our method were derived from BepiPred-2.0’s data set, which comprises of 11834 confirmed epitopes and 18722 confirmed non-epitopes. From those we removed all epitopes whose protein sequence of origin was shorter than 20 amino acid residues in length, resulting in our slightly modified version for BepiPred-2.0’s data set (*BepiPred-2.0 data set). Afterwards, we also removed all peptides contained in the training data sets of LBEEP, SVMTriP and LBtope, since the training and test sets of these methods were derived from IEDB, as was the case for BepiPred-2.0 data sets. The resulting data set was our first testing data set, Consensus Redundant (Consensus_R), which consists of 7675 epitopes and 15617 non-epitopes. From this data set we extracted a non-redundant data set by using CD-HIT – a method that allows the clustering of protein sequences based on their identity – with the sequence identity cutoff set at 0.6 (60%) and the rest of the settings left at their default values. Thus, our last data set was created, Consensus Non-Redundant (Consensus_NR), which contains 4286 epitopes and 5266 non-epitopes. Except for the original BepiPred-2.0 data set, all other data sets were then altered in order to create specialized data sets of fixed length peptides, using an extension and truncation methodology, as described in the main manuscript.
