## Supplementary File 2 for "Linear B-cell epitope prediction for *in silico* vaccine design: a performance review of methods available via command-line interface"

***Supplementary File 2	(Results and Discussion)***

**Linear B-cell epitope prediction for *in silico* vaccine design: a performance** **review of methods available via command-line interface**

**Kosmas A. Galanis^1^, Katerina C. Nastou^1, †^, Nikos C. Papandreou^1^, Georgios N. Petichakis^1^, Diomidis G. Pigis^1^ and Vassiliki A. Iconomidou^1,*^**

^1^Section of Cell Biology and Biophysics, Department of Biology, School of Sciences,

National and Kapodistrian University of Athens, Panepistimiopolis, Athens 15701, Greece

† Present Address: Novo Nordisk Foundation Center of Protein Research, University of Copenhagen, Blegdamsvej 3B, 2200 København, Denmark

Corresponding Author

*Associate Professor Vassiliki A. Iconomidou

Section of Cell Biology and Biophysics

Department of Biology

National and Kapodistrian University of Athens

Panepistimiopolis, Athens 15701, Greece

Performance of all predictors ­ except LBEEP ­ on Consensus_NR_exact

Further performance benchmarking was done on the Consensus_NR_exact data set, this time utilizing only the Consensus_NoLBEEP method, and all of the predictors except LBEEP. In the “per peptide” approach (Supplementary Table 5), a similar pattern with the Consensus_NR results emerges. The highest MCC score is achieved by BepiPred, with our Consensus_NoLBEEP method coming in second. BcePred is still characterized by high specificity, while ABCpred shows the best sensitivity. A slight shift in performance for the worst is observed by COBEpro and SVMTriP, where both methods score below 50% in accuracy and especially COBEpro, which also has a negative MCC.

**Supplementary Table 5. Performance of predictors in “per peptide” mode.** The methods are tested against the Consensus_NR_exact data set.

| *Predictor* | *SN%* | *SP%* | *ACC%* | *MCC* |
| --- | --- | --- | --- | --- |
| Consensus_NoLBEEP | 50.18 | 58.54 | 54.07 | 0.0873 |
| BcePred | 22.46 | 83.33 | 50.78 | 0.0726 |
| ABCpred | 67.93 | 37.08 | 53.59 | 0.0527 |
| LBtope | 46.2 | 59.58 | 52.42 | 0.0581 |
| BepiPred-1.0 | 52.72 | 56.88 | 54.65 | 0.0957 |
| COBEpro | 33.15 | 64.79 | 47.87 | -0.0216 |
| SVMTriP | 15.4 | 88.54 | 49.42 | 0.0574 |

The results obtained from the “per residue” approach (Supplementary Table 6) are quite similar with those reported on the Consensus_NR data set. Furthermore, the BepiPred and Consensus_RES classifiers still perform the best and COBEpro performs the poorest, with a negative MCC score of -0.0107. LBtope seems to perform a bit worse, with a lower MCC value, in contrast with BcePred which showed an increase in MCC value.

**Supplementary Table 6. Performance of “per residue” predictors.** The methods are tested against the Consensus_NR_exact data set.

| *Predictor* | *SN%* | *SP%* | *ACC%* | *MCC* |
| --- | --- | --- | --- | --- |
| Consensus_RES | 47.42 | 59.64 | 53.1 | 0.0709 |
| BcePred | 28.5 | 75.01 | 50.13 | 0.0395 |
| LBtope | 45.82 | 57.13 | 51.08 | 0.0296 |
| BepiPred-1.0 | 50.91 | 56.47 | 53.49 | 0.0737 |
| COBEpro | 37.09 | 61.86 | 48.61 | -0.0107 |

Performance of predictors on BepiPred-2.0’s data set at window size of 16

All predictors were also tested using a fixed window size of 16 residues, except for SVMTriP which unfortunately had no default threshold value set for the corresponding model. After initially obtaining poor results on the non-redundant data set, we also opted to use the entire BepiPred-2.0’s data set modified, so that all peptides had a fixed length of 16 residues, in order to ascertain the best possible performance for all methods.

Firstly, in the “per peptide” mode (Supplementary Table 7), the highest MCC score — by a wide margin — was achieved by LBtope, as well as the highest sensitivity, closely followed by BepiPred. The Consensus_noSVMTrip algorithm achieved the highest accuracy with 61.44%, but also exhibited an extremely low sensitivity and a specificity of nearly 100%, indicating that it rejected almost all sequences, perhaps as a result of a very high value in the consensus threshold. High specificity values were also observed for LBEEP, COBEpro and BcePred. Compared to the previous results on Consensus_NR with the window size of 20 amino acid residues, we observe a performance boost, especially in accuracy.

**Supplementary Table 7. Performance of predictors in “per peptide” mode. The methods are tested against BepiPred-2.0’s data set at fixed length of 16.**

| *Predictor* | *SN%* | *SP%* | *ACC%* | *MCC* |
| --- | --- | --- | --- | --- |
| Consensus_noSVMTrip | 1.39 | 99.41 | 61.44 | 0.0409 |
| BcePred | 27.37 | 75.48 | 56.81 | 0.0319 |
| ABCpred | 40.96 | 63.5 | 52.24 | 0.0459 |
| LBtope | 57.7 | 56.06 | 56.69 | 0.134 |
| BepiPred-1.0 | 53.98 | 53.86 | 53.91 | 0.0765 |
| COBEpro | 23.06 | 79.13 | 57.41 | 0.0259 |
| LBEEP | 23.14 | 79.73 | 57.81 | 0.0341 |

In the case of the “per residue” predictors (Supplementary Table 8), LBtope had once again the highest MCC, followed by our consensus method. Regarding the other metrics we see similar performance rankings with our previous test on the 20-residue peptides in MCC and accuracy. However, there is an overwhelming difference between the method’s sensitivity and specificity, rendering the method incapable of performing predictions in that window size.

**Supplementary Table 8. Performance of “per residue” predictors.** The methods are tested against BepiPred-2.0’s data set at fixed peptide length of 16.

| *Predictor* | *SN%* | *SP%* | *ACC%* | *MCC* |
| --- | --- | --- | --- | --- |
| Consensus_RES | 26.7 | 79.9 | 59.3 | 0.0769 |
| BcePred | 31.42 | 70.74 | 55.52 | 0.0231 |
| LBtope | 50.09 | 58.57 | 55.29 | 0.0849 |
| BepiPred-1.0 | 51.45 | 54.36 | 53.23 | 0.0566 |
| COBEpro | 36.19 | 64.86 | 53.75 | 0.0107 |

In short, changing the window size to 16 and including all available peptides of BepiPred-2.0’s data set, improved performance as expected, but led to a huge gap between – the previously balanced – sensitivity and specificity metrics. However, even with this advantage for the methods that were trained using peptides from IEDB (LBtope, LBEEP), the results remain unimpressive. Accuracy may have increased to ~58% for LBEEP and COBEpro, but their MCC values are still near zero indicating random classification. The performance improvement observed for LBEEP and LBtope is biased and probably associated with the inclusion of sequences already present in the training data sets of the two methods. On the other hand, ABCpred and COBEpro showed relatively improved accuracy without benefitting from the biased data set, which probably indicates their preference for shorter epitope segments.
