## Supplementary Table 1 for "Linear B-cell epitope prediction for *in silico* vaccine design: a performance review of methods available via command-line interface"

***Supplementary Table 1	(Introduction)***

**Linear B-cell epitope prediction for *in silico* vaccine design: a performance** **review of methods available via command-line interface**

**Kosmas A. Galanis^1^, Katerina C. Nastou^1, †^, Nikos C. Papandreou^1^, Georgios N. Petichakis^1^, Diomidis G. Pigis^1^ and Vassiliki A. Iconomidou^1,*^**

^1^Section of Cell Biology and Biophysics, Department of Biology, School of Sciences, National and Kapodistrian University of Athens, Panepistimiopolis, Athens 15701, Greece

† Present Address: Novo Nordisk Foundation Center of Protein Research, University of Copenhagen, Blegdamsvej 3B, 2200 København, Denmark

Corresponding Author

*Associate Professor Vassiliki A. Iconomidou

Section of Cell Biology and Biophysics

Department of Biology

National and Kapodistrian University of Athens

Panepistimiopolis, Athens 15701, Greece

In most cases, the algorithms that predict BCEs can either be sequence-based and/or structure-based. Most predictors utilize only data derived from the protein sequence of the antigen and thus are sequence-based, while structure-based predictors utilize only an antigen’s 3D structure. Furthermore, some hybrid methods employ both approaches for better predictive performance [1,2]. Historically, initial attempts at predicting epitopes made use of a single amino acid propensity scale, assigning each amino acid a numerical value, followed by a local averaging of these values along the peptide chain. The first method, implementing this approach, was published by Hopp and Woods [3] in 1981, and it utilized Levitt’s hydrophilicity scale [4]. Aside from hydrophilicity, which was utilized again in another scale by Parker et al. [5], other amino acid properties were explored in later methods, such as antigenicity [6], flexibility [7], surface accessibility [8], and turns [9]. The next wave of predictors built upon this development, when methods like PREDITOP [10], PEOPLE [11], BEPITOPE [12] and BcePred [13], combined multiple physicochemical properties. Although these methods represented the best attempts yet at predicting epitopes, Blythe and Flower [14] demonstrated that the performance of such methods was overstated. They did a thorough assessment of 484 amino acid propensity scales in combination with information on the location of epitopes for 50 known proteins and found that even the best possible combination of scales performed only slightly better than random [14]. In their work they also correctly suggested, that more advanced approaches for predicting linear B-cell epitopes needed to be developed, such as methods that employ artificial intelligence technology.

As anticipated, given the booming of available biological data, the entire next generation of methods utilized some form of machine learning models. One of the first such approaches was BepiPred [15], that combined a Hidden Markov Model (HMM) with an amino acid propensity scale. Additionally, other machine learning models were used in methods developed afterwards, including Neural Networks in ABCpred [16], a Naïve Bayes classifier in Epitopia [17] and Support Vector Machines (SVMs) in most of the recent predictors. SVM-based predictors dominated the machine learning approaches used in BCE prediction, each one differing from the other on feature selection, data set curation and SVM specific parameters (Supplementary Table 1). The BCPred [18] and FBCPred [19] methods published in 2008, predict fixed linear B-cell epitopes and flexible length linear B-cell epitopes respectively, utilizing SVM models with the subsequence kernel. The AAPPRED [20] method also utilizes SVM models trained on the frequency of Amino Acid Pairs (AAP), a scale first developed by Chen et al. [21]. Other notable approaches include: BayesB [22], LEPS [23] and BEORACLE [24]. A new machine learning approach that was developed in 2014, called EPMLR [25], utilizes multiple linear regression for epitope classification. Another recent novel approach is the DMN-LBE [26] method, that was developed using deep maxout networks, a type of deep neural network with a different activation layer called maxout. The DRREP [27] method was published in 2016, and it also utilizes deep neural network technology to extrapolate structural features related to epitopes from protein sequences. One of the latest additions is the second version of the BepiPred method, BepiPred-2.0 [2], that was developed in 2017. This method is based on a random forest algorithm and differs from its predecessor in that it was trained only on epitope data derived from crystal structures. Another promising algorithm is iBCE-EL [28], which is an ensemble learning framework combining Extremely Randomized Tree (ERT) and Gradient Boosting (GB) classifiers.

**Supplementary Table 1. Linear B-cell epitope predictors in chronological order, alongside a short description of their methodology, their current status and their web page.** After researching the relevant publications, we gathered up all the linear B-cell epitopes predictors we could find in this fairly complete, but not exhaustive catalogue. For every method we reference the source material to determine their methodology, which we have summed up for each predictor in a short description. For every predictor we also checked their availability status, as of writing this review, and categorized them regarding their general and current availability online as tools, as well as their obtainability as standalone software packages. In the last column, we provide the website links for each method, when available.

| *Predictor* | *Description* | *Status* | *Link* |
| --- | --- | --- | --- |
| Antigenic [6] | Physico-chemical propensity scales, occurrence of residues | Not currently available online | http://www.emboss.bioinformatics.nl/cgi-bin/emboss/antigenic |
| PEOPLE [11] | Physico-chemical propensity scales | Not available online | - |
| BEPITOPE [12] | Physico-chemical propensity scales | Freely available online | <http://bepitope.ibs.fr/> |
| BcePred [13] | Physico-chemical propensity scales | Freely available online and downloadable | <http://crdd.osdd.net/raghava/bcepred/index.html> |
| BepiPred-1.0 [15] | HMM & Parker hydrophilicity scale | Freely available online and downloadable | <http://www.cbs.dtu.dk/services/BepiPred-1.0/> |
| Söllner [29] | Physicochemical  Properties, MOE, KNN, Decision Tree | Not available online | - |
| Chen [21] | SVM & AAP | Not available online | - |
| ABCpred [16] | Neural networks (feed forward & reccurent) | Freely available online and downloadable | <http://crdd.osdd.net/raghava/abcpred/index.html> |
| BCPREDS [18,19] | SVM | Not currently available online | <http://ailab.ist.psu.edu/bcpred/> |
| AAPPred [20] | SVM & AAP | Freely available online and downloadable | <http://www.bioinf.ru/aappred/predict> |
| Epitopia [17] | ML algorithm trained to discern antigenic features | Freely available online and downloadable | <http://epitopia.tau.ac.il/index.html> |
| COBEpro [1] | SVM | Freely available online and downloadable | <http://scratch.proteomics.ics.uci.edu/> |
| BayesB [22] | SVM | Not currently available online | <http://immunopred.org/bayesb/index.html> |
| LEPS[23] | SVM & Physicochemical propensity scales & AAS | Not currently available online | <http://leps.cs.ntou.edu.tw/> |
| BEOracle [24] | SVM | Not available online | - |
| BEST [30] | SVM | Not available online | - |
| SVMTriP [31] | SVM | Freely available online and downloadable | <http://sysbio.unl.edu/SVMTriP/> |
| BEEPro [32] | SVM & Physicochemical propensity scales & PSSM | Not available online | - |
| LBtope [33] | SVM & Physicochemical propensity scales & AAP | Freely available online and downloadable | <http://crdd.osdd.net/raghava/lbtope/protein.php> |
| Random Forest [34] | Amino acid descriptors & Random Forest | Not currently available online | <http://sysbio.yznu.cn/Research/Epitopesprediction.aspx> |
| EPMLR [25] | Multiple Linear Regression | Not currently available online | <http://www.bioinfo.tsinghua.edu.cn/epitope/EPMLR/> |
| DMN-LBE [26] | Deep Maxout Networks | Not currently available online | <http://bioinfo.tsinghua.edu.cn/epitope/DMNLBE/> |
| LBEEP [35] | DDE - SVM | Freely available download | <https://github.com/brsaran/LBEEP> |
| APCpred [36] | APC & SVM | Not currently available online | <http://ccb.bmi.ac.cn/APCpred/> |
| DRREP [27] | Deep Ridge Neural Network | Not currently available online | <https://github.com/gsher1/DRREP> |
| BepiPred-2.0 [2] | Random forest algorithm trained on epitopes derived from crystal structures | Freely available online and downloadable | <http://www.cbs.dtu.dk/services/BepiPred/> |
| iBCE-EL [28] | Ensemble framework combining ERT & GB | Freely available online | <http://thegleelab.org/iBCE-EL/> |

**References**

1. Sweredoski MJ, Baldi P (2009) COBEpro: a novel system for predicting continuous B-cell epitopes. Protein Eng Des Sel 22 (3):113-120. doi:10.1093/protein/gzn075

2. Jespersen MC, Peters B, Nielsen M, Marcatili P (2017) BepiPred-2.0: improving sequence-based B-cell epitope prediction using conformational epitopes. Nucleic Acids Res 45 (W1):W24-W29. doi:10.1093/nar/gkx346

3. Hopp TP, Woods KR (1981) Prediction of protein antigenic determinants from amino acid sequences. Proc Natl Acad Sci U S A 78 (6):3824-3828

4. Levitt M (1976) A simplified representation of protein conformations for rapid simulation of protein folding. J Mol Biol 104 (1):59-107

5. Parker JM, Guo D, Hodges RS (1986) New hydrophilicity scale derived from high-performance liquid chromatography peptide retention data: correlation of predicted surface residues with antigenicity and X-ray-derived accessible sites. Biochemistry 25 (19):5425-5432

6. Kolaskar AS, Tongaonkar PC (1990) A semi-empirical method for prediction of antigenic determinants on protein antigens. FEBS Lett 276 (1-2):172-174

7. Karplus P, Schulz G (1985) Prediction of chain flexibility in proteins. Naturwissenschaften 72 (4):212-213

8. Emini EA, Hughes JV, Perlow DS, Boger J (1985) Induction of hepatitis A virus-neutralizing antibody by a virus-specific synthetic peptide. J Virol 55 (3):836-839

9. Pellequer JL, Westhof E, Van Regenmortel MH (1993) Correlation between the location of antigenic sites and the prediction of turns in proteins. Immunol Lett 36 (1):83-99

10. Pellequer JL, Westhof E (1993) PREDITOP: a program for antigenicity prediction. J Mol Graph 11 (3):204-210, 191-202

11. Alix AJ (1999) Predictive estimation of protein linear epitopes by using the program PEOPLE. Vaccine 18 (3-4):311-314

12. Odorico M, Pellequer JL (2003) BEPITOPE: predicting the location of continuous epitopes and patterns in proteins. J Mol Recognit 16 (1):20-22. doi:10.1002/jmr.602

13. Saha S, Raghava GPS BcePred: Prediction of Continuous B-Cell Epitopes in Antigenic Sequences Using Physico-chemical Properties. In, Berlin, Heidelberg, 2004. Artificial Immune Systems. Springer Berlin Heidelberg, pp 197-204

14. Blythe MJ, Flower DR (2005) Benchmarking B cell epitope prediction: underperformance of existing methods. Protein Sci 14 (1):246-248. doi:10.1110/ps.041059505

15. Larsen JE, Lund O, Nielsen M (2006) Improved method for predicting linear B-cell epitopes. Immunome Res 2:2. doi:10.1186/1745-7580-2-2

16. Saha S, Raghava GP (2006) Prediction of continuous B-cell epitopes in an antigen using recurrent neural network. Proteins 65 (1):40-48. doi:10.1002/prot.21078

17. Rubinstein ND, Mayrose I, Martz E, Pupko T (2009) Epitopia: a web-server for predicting B-cell epitopes. BMC Bioinformatics 10:287. doi:10.1186/1471-2105-10-287

18. El-Manzalawy Y, Dobbs D, Honavar V (2008) Predicting linear B-cell epitopes using string kernels. J Mol Recognit 21 (4):243-255. doi:10.1002/jmr.893

19. El-Manzalawy Y, Dobbs D, Honavar V (2008) Predicting flexible length linear B-cell epitopes. Comput Syst Bioinformatics Conf 7:121-132

20. Davydov YI, Tonevitsky AG (2009) Prediction of linear B-cell epitopes. Molecular Biology 43 (1):150-158. doi:10.1134/s0026893309010208

21. Chen J, Liu H, Yang J, Chou KC (2007) Prediction of linear B-cell epitopes using amino acid pair antigenicity scale. Amino Acids 33 (3):423-428. doi:10.1007/s00726-006-0485-9

22. Wee LJ, Simarmata D, Kam YW, Ng LF, Tong JC (2010) SVM-based prediction of linear B-cell epitopes using Bayes Feature Extraction. BMC Genomics 11 Suppl 4:S21. doi:10.1186/1471-2164-11-S4-S21

23. Wang HW, Lin YC, Pai TW, Chang HT (2011) Prediction of B-cell linear epitopes with a combination of support vector machine classification and amino acid propensity identification. J Biomed Biotechnol 2011:432830. doi:10.1155/2011/432830

24. Wang Y, Wu W, Negre NN, White KP, Li C, Shah PK (2011) Determinants of antigenicity and specificity in immune response for protein sequences. BMC Bioinformatics 12:251. doi:10.1186/1471-2105-12-251

25. Lian Y, Ge M, Pan XM (2014) EPMLR: sequence-based linear B-cell epitope prediction method using multiple linear regression. BMC Bioinformatics 15:414. doi:10.1186/s12859-014-0414-y

26. Lian Y, Huang ZC, Ge M, Pan XM (2015) An Improved Method for Predicting Linear B-cell Epitope Using Deep Maxout Networks. Biomed Environ Sci 28 (6):460-463. doi:10.3967/bes2015.065

27. Sher G, Zhi D, Zhang S (2017) DRREP: deep ridge regressed epitope predictor. BMC Genomics 18 (Suppl 6):676. doi:10.1186/s12864-017-4024-8

28. Manavalan B, Govindaraj RG, Shin TH, Kim MO, Lee G (2018) iBCE-EL: A New Ensemble Learning Framework for Improved Linear B-Cell Epitope Prediction. Front Immunol 9:1695. doi:10.3389/fimmu.2018.01695

29. Sollner J, Mayer B (2006) Machine learning approaches for prediction of linear B-cell epitopes on proteins. J Mol Recognit 19 (3):200-208. doi:10.1002/jmr.771

30. Gao J, Faraggi E, Zhou Y, Ruan J, Kurgan L (2012) BEST: improved prediction of B-cell epitopes from antigen sequences. PloS one 7 (6):e40104. doi:10.1371/journal.pone.0040104

31. Yao B, Zhang L, Liang S, Zhang C (2012) SVMTriP: A Method to Predict Antigenic Epitopes Using Support Vector Machine to Integrate Tri-Peptide Similarity and Propensity. PLOS ONE 7 (9):e45152. doi:10.1371/journal.pone.0045152

32. Lin SY, Cheng CW, Su EC (2013) Prediction of B-cell epitopes using evolutionary information and propensity scales. BMC bioinformatics 14 Suppl 2:S10

33. Singh H, Ansari HR, Raghava GP (2013) Improved method for linear B-cell epitope prediction using antigen's primary sequence. PLoS One 8 (5):e62216. doi:10.1371/journal.pone.0062216

34. Huang JH, Wen M, Tang LJ, Xie HL, Fu L, Liang YZ, Lu HM (2014) Using random forest to classify linear B-cell epitopes based on amino acid properties and molecular features. Biochimie 103:1-6. doi:10.1016/j.biochi.2014.03.016

35. Saravanan V, Gautham N (2015) Harnessing Computational Biology for Exact Linear B-Cell Epitope Prediction: A Novel Amino Acid Composition-Based Feature Descriptor. OMICS 19 (10):648-658. doi:10.1089/omi.2015.0095

36. Shen W, Cao Y, Cha L, Zhang X, Ying X, Zhang W, Ge K, Li W, Zhong L (2015) Predicting linear B-cell epitopes using amino acid anchoring pair composition. BioData Min 8:14. doi:10.1186/s13040-015-0047-3
