## Supplementary Table 2 for "Linear B-cell epitope prediction for *in silico* vaccine design: a performance review of methods available via command-line interface"

***Supplementary Table 2	(Materials and Methods)***

**Linear B-cell epitope prediction for *in silico* vaccine design: a performance** **review of methods available via command-line interface**

**Kosmas A. Galanis^1^, Katerina C. Nastou^1, †^, Nikos C. Papandreou^1^, Georgios N. Petichakis^1^, Diomidis G. Pigis^1^ and Vassiliki A. Iconomidou^1,*^**

^1^Section of Cell Biology and Biophysics, Department of Biology, School of Sciences,

National and Kapodistrian University of Athens, Panepistimiopolis, Athens 15701, Greece

† Present Address: Novo Nordisk Foundation Center of Protein Research, University of Copenhagen, Blegdamsvej 3B, 2200 København, Denmark

Corresponding Author

*Associate Professor Vassiliki A. Iconomidou

Section of Cell Biology and Biophysics

Department of Biology

National and Kapodistrian University of Athens

Panepistimiopolis, Athens 15701, Greece

Selection of suitable linear B-cell epitope predictors

BcePred was published in 2004 by Raghava *et al*. [1], and is based on a plethora of physicochemical propensity scales utilizing amino acid properties, such as hydrophilicity and antigenicity, either individually or in combination. Moreover, it achieved a reported 56% sensitivity, 61% specificity and its highest accuracy of 58.70%, on a data set obtained from the database Bcipep [2], using a combination of flexibility, hydrophilicity, polarity and surface accessibility propensity scales.

BepiPred was developed in 2006 by Lund *et al*.[3], and it is the first ever method that utilizes an HMM. The HMM was trained using a data set derived from the database Antijen [4] and the Pellequer data set [5], and was then combined with Parker’s hydrophilicity scale, resulting in the BepiPred method. This method managed to achieve an Area Under Curve (AUC) of the Receiver Operating Characteristic (ROC) curve of 0.671 ± 0.013 on the Pellequer data set.

ABCpred was created in 2006 [6], again by the Raghava group and it was the first test case of a more sophisticated machine learning model. It is based on a Recurrent Neural Network (RNN) that was trained using a variety of different window sizes and hidden units. The window sizes that were tested, were 10, 12, 14, 16, 18 and 20. Thus six models were developed in total, with the window size of 16 amino acid residues achieving the highest accuracy of 65.93% and a Matthews Correlation Coefficient (MCC) of 0.3187, after five-fold cross-validation on a data set derived from Bcipep [2].

COBEpro was published in 2009 by Baldi *et* al. [7] at the University of California. This method utilizes a novel two-step system for the prediction of both linear and discontinuous B-cell epitopes. Firstly, it utilizes an SVM model to assign an epitopic propensity score to fragments within the given peptide sequence. Additionally, COBEpro is able to incorporate into the SVM model the provided or predicted secondary structure and solvent accessibility of the given sequence, that are predicted by SSpro [8] and ACCpro [9] respectively. During the second stage, the method calculates an epitopic propensity score for each amino acid, based on the previous scores assigned by the model in the first stage. Among others, this predictor was tested on the fragmented version of Chen’s [10] data set, achieving an AUC of 0.829 and an accuracy of 78%.

SVMTriP was developed in 2012 [11] and it is an application of an SVM model that employs tri-peptide similarity calculated through the Blosum62 matrix in combination with amino acid propensity scales. Its prediction suite comes with six different models corresponding to window sizes of 10, 12, 14, 16, 18 and 20 of which the 20 amino acid residue model performed the best with a reported 80.10% sensitivity and 55.20% precision on a data set gathered from the Immune Epitope Data Base (IEDB) [12].

LBtope was the most recent effort, out of our selected predictors,on epitope prediction published by Raghava’s lab in 2013. This method uses, among other previously used types of features, a modified AAP profile from Chen’s method [10]. These profiles are used to convert the input sequence into numerical features that are then used as input for an SVM model that predicts epitopes. LBtope was trained and tested on a data set collected from IEDB, which comprised of experimentally verified epitopes and non-epitopes, in contrast to previous methods that used random peptides as non-epitopes. Its reported performance on different data sets varied significantly, with an accuracy ranging from 51.57% to 85.74%.

LBEEP was developed in 2015 by Saravan *et al*. [13] from the University of Madras in India. In this work, a novel amino acid feature descriptor called Dipeptide Deviation from Expected Mean (DDE) was developed, in an attempt to distinguish linear epitopes from non-epitopes. This new descriptor was then implemented with both SVM and AdaBoost-Random Forest machine learning techniques. The data set used to train this method was constructed by using only exact epitopes, instead of epitope containing regions, which have been used as training material in the past, making LBEEP a pioneer method in that respect. The exact epitopes used for training were isolated from IEDB [14] and are 5 to 15 amino acid residues long and thus LBEEP is better suited for predictions in that range. During testing, LBEEP achieved an accuracy between 61% and 73%, after five-fold cross-validation, on a data set derived also from IEDB.

Development of the consensus method

**Supplementary Table 2. Input window sizes and prediction approach of each method.** The classification of query proteins as epitopes can generally be performed in either a “per residue” or a “per peptide” basis. In the “per residue” methods each separate residue of a protein is assigned an antigenicity score, while in the “per peptide” methods, a prediction is limited within fixed windows sizes.

| *Predictor* | *Prediction* | *Window Size* |
| --- | --- | --- |
| ABCpred | Per peptide | 10, 12, 14, 16, 18, 20 |
| SVMTriP | Per peptide | 10, 12, 14, 16, 18, 20 |
| LBEEP | Per peptide | 5 - 15 |
| BcePred | Per residue | - |
| BepiPred-1.0 | Per residue | - |
| COBEpro | Per residue | - |
| LBtope | Per residue | - |

The methods that predict per peptide, ABCpred and SVMTriP, use predetermined fixed window sizes. Thus, it was necessary to choose a window size where these methods would operate sufficiently well, both in individual testing and as part of the consensus classifier. The window size chosen for these methods after initial testing was that of 20 residues. The main reasons were the better reported performance of SVMTriP at that window size and the lack of any default threshold values for the rest of the models in the documentation. As far as ABCpred is concerned, the performance penalty of selecting a window size of 20 instead of the reported best of 16 residues was minor. It should also be noted that initial testing for LBEEP at a window size of 20 was experimental, since the method was trained using only epitopes of lengths between 5 and 15, and thus any results outside that range were unreliable. Moreover, using the ‘confirmed model’, as suggested by the creators of the method in the GitHub repository of LBEEP, returned worse results than the default ‘balanced’ model, and so we opted to use the latter (Supplementary Table 4B).

Once a window size of 20 was selected for the “per peptide” methods, an effective strategy had to be formulated where the two different categories of output would produce a single consensus result. The solution was a consensus voting system that classifies a residue as belonging to an epitope when a predetermined threshold of votes has been achieved. When a “per residue” method classifies a residue of the query sequence as “epitopic” it counts as one positive vote, while when a “per peptide” method classifies a fragment of a protein as an epitope each amino acid of that peptide is classified as “epitopic”. So, when the sum of positive votes for a given position of a query sequence surpasses the threshold of the consensus classifier, that residue is marked as part of an epitope. The consensus threshold chosen, after testing, is defined as the hit overlap of at least half out of “n” selected methods, where “n” is the number of methods embedded in the algorithm [15]. The consensus method accepts protein sequences, of a length of 20 amino acid residues or higher, in FASTA format as input. The workflow of the consensus method is shown in Supplementary File 2.

For testing purposes, a slightly different architecture of the consensus method was implemented, that specialized in rapid consensus output on our fixed length data sets (Figure 1). All methods – including the consensus – were mainly tested on a data set consisting of peptides with a length of 20. To resolve this issue, two parallel approaches were explored. In the first approach, all methods were included, and each method predicted whether an entire peptide is an epitope or not. However, in order for the results between the “per peptide” and “per residue” methods to be comparable, since only “per peptide” methods classify protein fragments, it was accepted that when “per residue” methods have predicted half or more of a peptide’s fragments as “epitopic”, then the whole peptide too is a predicted epitope. Such caveats are generally found in other forms of predictors of biological nature [16,17], and thus were chosen in our evaluation approach, as well. In the second approach, only “per residue” methods were included, and the consensus result was simply, a combination of only those predictions.

**References**

1. Saha S, Raghava GPS BcePred: Prediction of Continuous B-Cell Epitopes in Antigenic Sequences Using Physico-chemical Properties. In, Berlin, Heidelberg, 2004. Artificial Immune Systems. Springer Berlin Heidelberg, pp 197-204

2. Saha S, Bhasin M, Raghava GP (2005) Bcipep: a database of B-cell epitopes. BMC Genomics 6:79. doi:10.1186/1471-2164-6-79

3. Larsen JE, Lund O, Nielsen M (2006) Improved method for predicting linear B-cell epitopes. Immunome Res 2:2. doi:10.1186/1745-7580-2-2

4. Toseland CP, Clayton DJ, McSparron H, Hemsley SL, Blythe MJ, Paine K, Doytchinova IA, Guan P, Hattotuwagama CK, Flower DR (2005) AntiJen: a quantitative immunology database integrating functional, thermodynamic, kinetic, biophysical, and cellular data. Immunome Res 1 (1):4. doi:10.1186/1745-7580-1-4

5. Pellequer JL, Westhof E, Van Regenmortel MH (1993) Correlation between the location of antigenic sites and the prediction of turns in proteins. Immunol Lett 36 (1):83-99

6. Saha S, Raghava GP (2006) Prediction of continuous B-cell epitopes in an antigen using recurrent neural network. Proteins 65 (1):40-48. doi:10.1002/prot.21078

7. Sweredoski MJ, Baldi P (2009) COBEpro: a novel system for predicting continuous B-cell epitopes. Protein Eng Des Sel 22 (3):113-120. doi:10.1093/protein/gzn075

8. Pollastri G, Baldi P, Fariselli P, Casadio R (2002) Prediction of coordination number and relative solvent accessibility in proteins. Proteins 47 (2):142-153

9. Cheng J, Randall AZ, Sweredoski MJ, Baldi P (2005) SCRATCH: a protein structure and structural feature prediction server. Nucleic Acids Res 33 (Web Server issue):W72-76. doi:10.1093/nar/gki396

10. Chen J, Liu H, Yang J, Chou KC (2007) Prediction of linear B-cell epitopes using amino acid pair antigenicity scale. Amino Acids 33 (3):423-428. doi:10.1007/s00726-006-0485-9

11. Yao B, Zhang L, Liang S, Zhang C (2012) SVMTriP: A Method to Predict Antigenic Epitopes Using Support Vector Machine to Integrate Tri-Peptide Similarity and Propensity. PLOS ONE 7 (9):e45152. doi:10.1371/journal.pone.0045152

12. Vita R, Zarebski L, Greenbaum JA, Emami H, Hoof I, Salimi N, Damle R, Sette A, Peters B (2010) The immune epitope database 2.0. Nucleic Acids Res 38 (Database issue):D854-862. doi:10.1093/nar/gkp1004

13. Saravanan V, Gautham N (2015) Harnessing Computational Biology for Exact Linear B-Cell Epitope Prediction: A Novel Amino Acid Composition-Based Feature Descriptor. OMICS 19 (10):648-658. doi:10.1089/omi.2015.0095

14. Vita R, Overton JA, Greenbaum JA, Ponomarenko J, Clark JD, Cantrell JR, Wheeler DK, Gabbard JL, Hix D, Sette A, Peters B (2015) The immune epitope database (IEDB) 3.0. Nucleic Acids Res 43 (Database issue):D405-412. doi:10.1093/nar/gku938

15. Frousios KK, Iconomidou VA, Karletidi CM, Hamodrakas SJ (2009) Amyloidogenic determinants are usually not buried. BMC Struct Biol 9:44. doi:10.1186/1472-6807-9-44

16. Peters C, Tsirigos KD, Shu N, Elofsson A (2016) Improved topology prediction using the terminal hydrophobic helices rule. Bioinformatics 32 (8):1158-1162. doi:10.1093/bioinformatics/btv709

17. Kall L, Krogh A, Sonnhammer EL (2004) A combined transmembrane topology and signal peptide prediction method. J Mol Biol 338 (5):1027-1036. doi:10.1016/j.jmb.2004.03.016
